# Establishing remote communication between human operator and engineered intestinal microbe via bio-electronic optical language

**DOI:** 10.1101/2024.05.24.595785

**Authors:** Xinyu Zhang, Zhijie Feng, Lianyue Li, Haoyan Yang, Pengxiu Dai, Chao Yang, Hongxiang Li, Dawei Sun, Hanxin Wang, Huimin Xue, Yaxin Wang, Xinyu Liu, Mingshan Li, Shenjunjie Lu, Meihui Cui, Xiaoyan Ma, Jing Liu, Yanyan Shi, Taofeng Du, Duo Liu, Hanjie Wang

## Abstract

Synthetic biology enables microbes to be engineered to detect biomarkers and deliver therapies *in situ*, offering advantages such as self-replication and localized response and treatment. However, the engineered microbes in the intestine remains a “black box”, the status of microbial functionality is inaccessible, and the activation of microbial functions is uncontrollable. There is a demand of a “language” between microbes and human for real-time and *in-situ* monitoring and controlling engineered microbes in the intestine. Light, due to its advantages of specificity and accuracy in the body, was chosen as the language to establish an interactive communication path between human and the microbes with the features of sophisticated monitoring and controlling. For effectively remote operation of this “language” in the intestine, we introduced an electronic capsule as the intermedia, translating information and instructions understandable for human operator into bacterial regulatory and detectable signals *in vivo*. In this paper, the electronic capsule and the engineered *Escherichia coli* Nissle 1917 (EcN) strains were designed in a collaborative manner. An EcN-to-capsule bioluminescent signal transmission was established and testified as the language to report the status of engineered EcN, especially the response of EcN to intestinal environment. Similarly, another capsule-to-EcN optogenetic signal transmission was established and testified as the language to control the function of engineered EcN, especially the secretion of therapeutic substances. On this basis, bidirectional communication between human operator and intestinal microbes was achieved via bidirectional bio-electronic optical language. The communication was established to monitor and regulate engineered EcN community in the intestine in 50-90 kg live pigs. The *in vitro* and *in vivo* estimations demonstrated the capabilities of monitoring and regulating the engineered microbes, offering a remotely controllable language-based communication between human operator and internal intestinal microbes.

## Introduction

Synthetic biology enables bacteria to be engineered for diagnosis and treatment through responding to biomarkers and expressing therapeutic genes in the intestine^1,2^. Engineered microbes have exhibited potentiality in treating diseases such as gastrointestinal diseases^3,4^, cancers^5^, metabolic syndromes^6–8^, *etc*. However, the engineered microbes in the intestine remains a “black box”, the statuses of engineered microbes are inaccessible, and the function of engineered microbes is uncontrollable. Due to the greater tissue depth and the complicated physiological gut environment, it is difficult and indirect to monitor and control engineered microbes *in-situ* and real-time. The challenges in monitoring and controlling engineered microbes have led to a poor understanding of their functional process in the intestine.

Therefore, there is a demand of a “language” between microbes and human to real-time and *in-situ* monitor and control engineered microbes in the intestine. Ingestible devices such as electronic capsules have been endowed with intestinal *in-situ* functions such as deep endoscopy^9^, sample collection^10^, signal detection^11,12^, gut stimulation^13^, and drug delivery^14^. Recently, bio-electronic capsules were developed that packaging engineered microbes as whole-cell sensors in the capsule for detecting bioluminescence that reflecting intestinal markers^15,16^. These progresses have indicated the convenience of detection and operation of the intestine using ingestible electronic capsules, and the possibility for using electronic capsules as a “encoding and decoding processor” to transfer signals and instructions from the human operator into bacterial regulatable and detectable signals *in vivo*.

For the purpose of monitoring and regulating intestinal engineered microbes, the bacterial strains and the capsule should be designed as collaborative, with efficient bi-directional communication. Light, as a bridge connecting microscopic biological processes and macroscopic experimental methods, have been extensively applied in *in-vivo* researches^17,18^, due to its advantages in extremely low crosstalk, spatiotemporal precision, and orthogonality. In this research, optical signals are also a suitable candidate as the “language” for bi-directional communication between engineered bacterial strains and the electronic capsule. Specifically, engineered bacterial strains express luciferase^19^ to be received by photoelectric sensor of the capsule, or respond to optogenetic instuctions^20^ from LEDs of the capsule.

In this article, we report a strategy that developing an optical language between engineered microbes and electronic capsules, and establishing a bridge of information for the bi-directional communication between human and engineered microbes *in situ* (**Fig. 1**). In this article, *Escherichia coli* Nissle 1917 (EcN) was chosen as the chassis to be engineered with the ability to receive optogenetic “language”, send bioluminescent “language”, and function in the host’s intestine. An electronic capsule was developed as an intermedia to encoding and decoding information by remotely sending information to and receiving instructions from operators, as well as emitting optogenetic signals to and detecting bioluminescent from bacteria. The bidirectional optical communication between microbes and the capsule demonstrated the potential in application was examined both *in vitro* and *in vivo* on live pigs. The bi-directional optical communication provides an *in-situ* and real-time method to monitor and regulate intestinal engineered microbes remotely, and shows potential to realize the closed-loop diagnosis and treatment under the supervision and control of operators.

**Figure 1.**
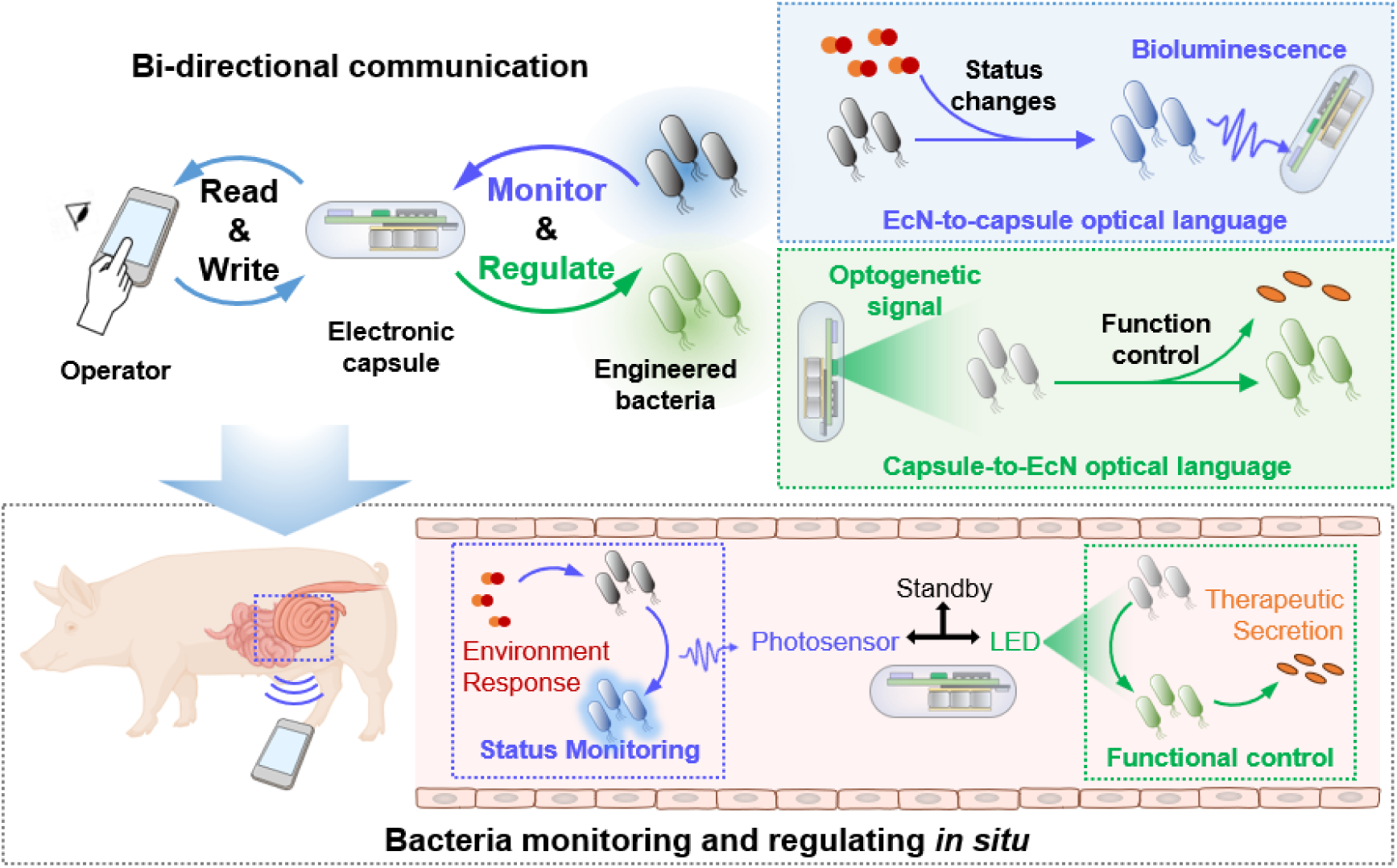
Overview of the optical communication for engineered bacteria monitoring and controlling. The electronic capsule and engineered EcN strains were designed in a collaborative manner for bi-directional optical communication as a kind of language. The strategy was based on the EcN-to-capsule bioluminescent signal transmission for bacterial status monitoring, and the capsule-to-EcN optogenetic signal transmission for bacterial function regulation. When applied *in vivo*, the capsule was remotely controllable through an APP operated by human beings to monitor and control the engineered EcNs in the intestine of live pigs.

## Result

### The EcN-to-capsule optical signal transmission for microbial status monitoring

For the optical language communication between the engineered EcN and the capsule, the patterns of optical signal transmission would determine the capacity of monitoring the microbial state and regulating the microbial functionality. The *lux-ABCDE* coding luciferase module^19^ equipped bacteria with the ability of simultaneously synthesizing luciferase and its cellular reacting substrate fatty aldehyde, allowing the engineered EcN to autonomously emit bioluminescence as optical language signals that could be transmitted to the electronic capsule. In our strategy, engineered EcN expressed luciferase in a constitutive or inducible manner to respond to particular changes of the engineered EcN status in intestine, such as the changes of cell density and environmental molecular concentration (**Fig 2A**). As an intermediate of signal and information transmitting between human operator and the engineered bacteria, the capsule could play a role of synchronized microbial -electronic language translator. This capsule was consisted of a photosensor to detect EcN bioluminescence, a LED to stimulate opto-effector bio-module in EcN, and a wireless Bluetooth module (**Fig 2B**). The section diameter of the electronic capsule was about 14 mm and the long-end length was about 33 mm.

**Figure 2.**
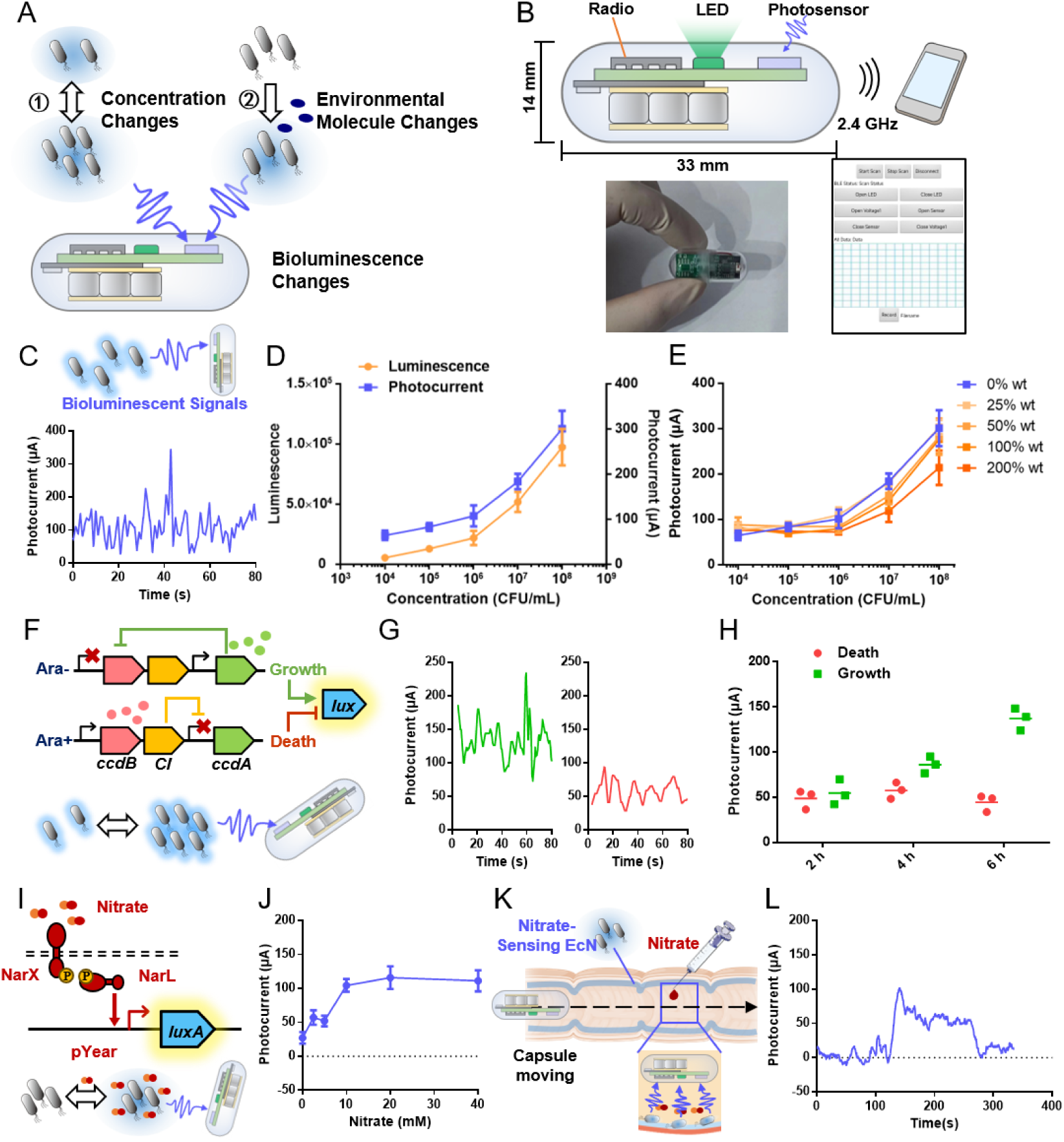
EcN-to-capsule bioluminescent signal transmission for microbial status monitoring. (A) Schematic diagram of the EcN-to-capsule bioluminescent signal transmission as the language to report the microbial status changes, including 1) cellular concentration changes and 2) environmental molecular changes. **(B)** Schematic diagram of the components and structure of the capsule and the APP panel for users to remotely control the capsule. **(C)** The bioluminescent signal transmission from constitutive *Lux-ABCDE* expressing EcN to capsule. **(D)** The bioluminescent values of different concentrations of constitutive Lux expressing EcN, detected by the microplate reader (Left Y-axis) and by the capsule (Right Y-axis). **(E)** The bioluminescent language received by the capsule that received from different concentrations of constitutive *Lux-ABCDE* expressing EcN, across transmission medium of simulated chyme composed of saline mixed with solid feed. **(F)** The EcN-to-capsule bioluminescent language reporting bacterial concentration under the control of a survival-death genetic circuit. **(G)** Representative bioluminescent signals monitored by the capsule of un-induced bacterial growth and induced bacterial death after 6 hours. **(H)** The mean values of bioluminescence monitored by the capsule from un-induced bacterial growth and induced bacterial death. **(I)** The EcN-to-capsule bioluminescent language reporting the response of environmental nitrate concentration. **(J)** The mean values of bioluminescence detected by the capsule indicating the nitrate-sensing EcN response to different concentrations of nitrate for 2 hours. **(K)** The *in vitro* simulation experiment in which the capsule monitored nitrate distribution while passing through a segment of intestinal tissue evenly distributed nitrate-sensing EcN. **(L)** The bioluminescent signal monitored by the capsule while the capsule passing through the segment of intestinal tissue. (Data are presented as the mean ± SD. n=3. * 0.01<p<0.05, ** 0.001<p<0.01, *** p<0.001.)

When the inside photosensor was turned on, the capsule could record the bioluminescent signals emitted from the luciferase-expressing EcN. Under dark conditions, the capsule could receive weak bioluminescent signals from constitutive luciferase-expressing EcN strains (**Fig 2C**). Theoretically, in an intestinal environment with relatively stable distribution of engineered EcNs, the EcN-to-capsule bioluminescence signaling would be mainly influenced by bacterial concentration and unexpected light-absorbing media such as chyme in the intestine. Comparing the bioluminescent intensity of constitutive luciferase-expressing EcN measured by the capsule against the value measured by standard microplate reader devices, it was demonstrated that the bacterial bioluminescence signal transmission recorded by the capsule was dose-accurate (**Fig 2D**), exhibiting a linear correlation with standard testing methods (R^2^ = 0.94) (**Fig S1**). It was noticed that the raw data of the photocurrent value exhibited drastic rapid data fluctuations. To cope with this situation, a sliding average filtering code was introduced for smooth output (**Fig S2**). This algorithm would not affect the arithmetic mean value of the data. Next, we simulated chyme using different concentrations of feed mixed in saline, the bioluminescent signals of constitutive luciferase-expressing EcN were weakened when the concentration of EcN was low or the concentration of feed was high (**Fig 2E**). On the contrary, when the EcN population density reached higher 10^7^ or even 10^8^, the effect of feed was relatively weak, suggesting that the sufficient bacterial concentration applied *in vivo* was the basis for monitoring engineered bacteria.

To demonstrate the application potential of the constitutive EcN-to-capsule bioluminescent signal transmission, a scenario was devised to monitor surviving bacterial density changes under artificial regulation (**Fig 2F**). Wherein, the engineered EcN was equipped with a survival-death switch genetic circuit for artificial controlling of cell growth or demise by using L-arabinose as an inducer (referred as survival-death switch EcN strain). The experiment result showed that the engineered bacterial density changes could be successfully monitored by the capsule in the form of bioluminescence signals (**Fig 2G**). When monitoring survival-death switch EcN strain with the capsule, the surviving group without arabinose-induction showed rapid growth and significant enhancement of EcN-to-capsule bioluminescent signal, while the arabinose-induced death group showed much slower bacterial growth and reversed concentration, transmitting much lower intensity of bioluminescence signals to the capsule (**Fig 2H and S2**).

Another scenario to evaluate the application potential of the EcN-to-capsule signal transmission was devised that using environmental signal sensing EcN to reflect the molecular changes. L-arabinose was taken as a pattern molecule to be sensing firstly. An arabinose-responsive bioluminescent strain (referring as ara-sensing EcN) was constructed in order to examine the EcN-to-capsule signal transmission flux indicating the concentration of arabinose changes (**Fig S3A**). Ara-sensing EcN presented almost no leakage of reporter gene expression when no arabinose was added, but instead showed rapidly strong reporter gene expression after sensing arabinose both *in vitro* and in live mouse intestine *in vivo* (**Fig S3B-D**). In further experiment, the photocurrent values of the capsule clearly indicated the EcN-to-capsule signals of ara-sensing EcN which being induced by different concentration of arabinose (**Fig S3E and F**). Uneven distribution of molecules in the intestine was simulated with an intestinal tissue *in vitro*.

Ara-sensing EcN was evenly added inside the inner wall of the intestinal tissue, and then 100 μL 10 mM arabinose solution was injected inside the inner wall and incubated for 2 hours. When operating the capsule to pass through the intestinal tissue, the bioluminescence from evenly distributed ara-sensing EcN significantly increased only in areas with higher concentration of arabinose, and was easily distinguished by the capsule during its passage through the intestine (**Fig S3G-I**).

In order to extend the above scenario to more practical applications for diagnosing intestinal health, a nitrate-responsive bioluminescent strain (referring as nitrate-sensing EcN strain) was constructed to transmit bioluminescence signals that indicate nitrate concentration around the capsule (**Fig 2I**). In the intestinal physiological conditions, nitrate is generated from inducible nitric oxide synthase (iNOS) that highly expressed in the inflammatory intestinal epithelial tissues, thus becomes a biomarker of intestinal inflammatory. Although there was a background leakage expression in the absence of nitrate, nitrate-sensing EcN quickly responded to different concentrations of nitrate within 2 hours (**Fig S4A-C**). Nitrate-sensing EcN also showed obvious differences of bioluminescence intensities in the intestines of mice exposed to exogenous 10 mM nitrate (**Fig S4D**). Detecting through EcN-to-capsule signal transmission, nitrate with a concentration equal to or greater than 10 mM could be significantly measured (**Fig 2J, S4E and S4F**). Uneven distribution of nitrate was also simulated with an intestinal tissue *in vitro* by even addition of nitrate-sensing EcN and particularly local injection of 100 μL 10 mM nitrate solution (**Fig 2K and S4G**). When operating the capsule to pass through the intestinal tissue, the local higher concentration of nitrate could be easily recognized by the capsule during it passing through the intestine (**Fig 2L**).

### The capsule-to-EcN optical signal transmission for microbial function control

In our strategy, light signals as the optical language transmitted from capsule to the engineered EcN carried the information of activating the optogenetic circuit function in the strain. Optimized CcaS-CcaR optogenetic two-component system was reported to achieve the strongest induction factor among all of the two-component optogenetic tools^21^. The green light optogenetic tool based on CcaS-CcaR has been applied to control the production of colanic acid in engineered bacteria in the gut of *Caenorhabditis elegans*, thus achieving light-controlled promotion of longevity^22^. In this study, the CcaS-CcaR optogenetic circuit was introduced to construct an opto-effector EcN strain. The optogenetic function of this strain would be motivated by the green light emitted from the LEDs equipped in the electron capsule (**Fig 3A and Fig S5A**).

**Figure 3.**
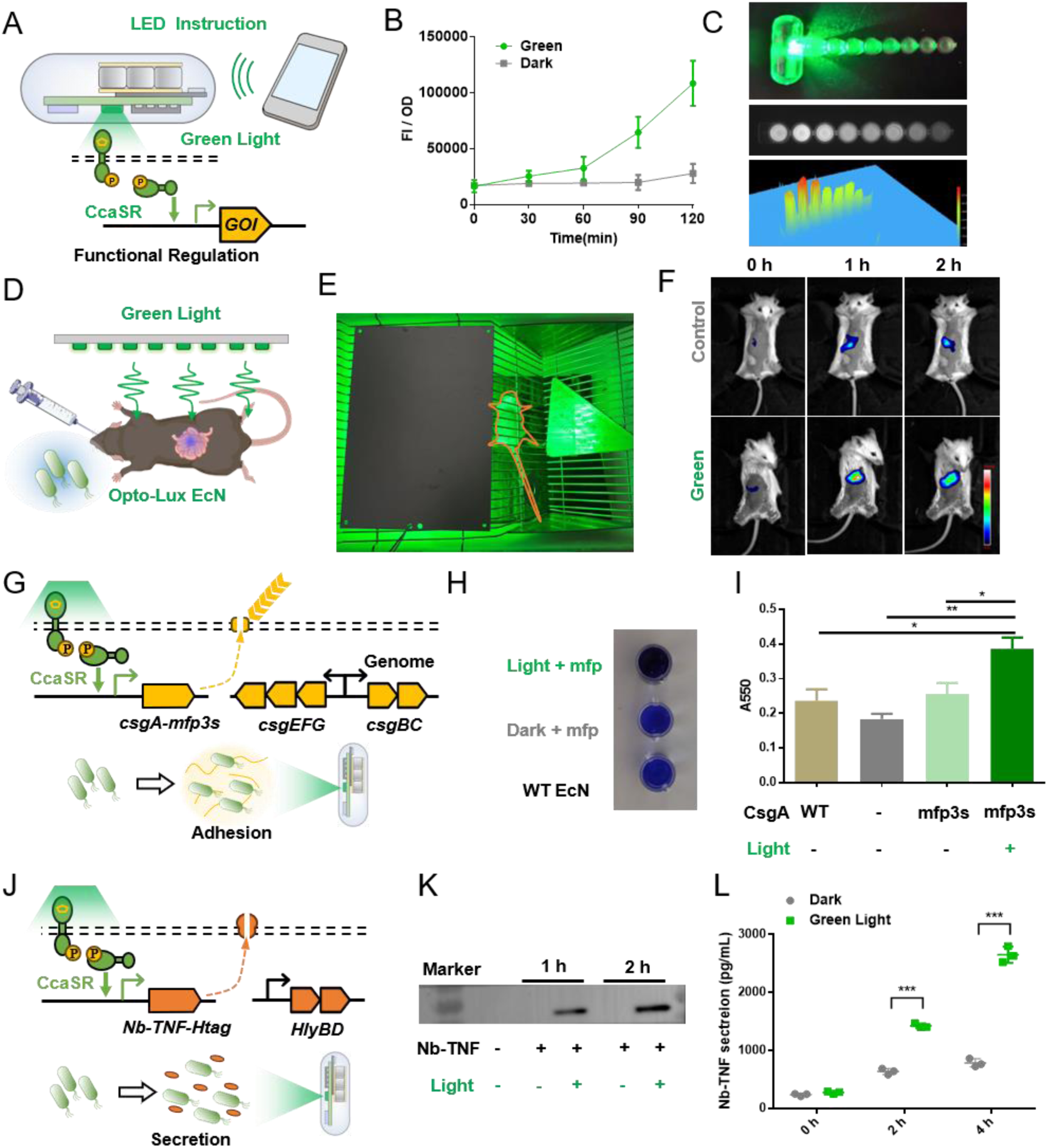
Capsule-to-EcN optogenetic signal transmission for microbial function control. (A) Schematic diagram of the capsule-to-EcN optogenetic signal transmission as the language to regulate the expression of gene of interest (GOI). **(B)** Fluorescence intensity of sfGFP induced by capsule-to-EcN green light within 2 hours. **(C)** The special performance of capsule-to-EcN optogenetic regulation. Using opto-effector EcN activated in microwells to explore the spatial characteristics of capsule-regulated engineered bacteria (top). Qualitative (middle) and quantitative (bottom) detection of sfGFP expression in each pore by fluorescence imaging after 2 hours of irradiation. **(D)** Schematic diagram of green light irradiation strategy for opto-Lux EcN-gavage mice. **(E)** Experimental photograph of green light irradiation strategy for opto-Lux EcN-gavage mice. **(F)** Luminescence mode imaging of opto-Lux EcN-gavage mice with and without green light irradiation. **(G)** Schematic diagram of capsule-to-EcN optogenetic regulation of the formation of the adhesive CsgA-mfp3s matrix by opto-adhesion EcN strain. **(H)** Validation of the enhanced adhesion effect of green light-induced opto-adhesion EcN strain through crystal violet staining absorbance at 550 nm. **(I)** Photograph of bacterial adhesion under capsule-to-EcN green light control. **(J)** Schematic diagram of capsule-to-EcN optogenetic regulation of the secretion of Nb-TNF. **(K)** Verification of Nb-TNF secretion in the supernatant induced by capsule-to-EcN light for 2 hours using WB. **(L)** Quantification of Nb-TNF secretion in the supernatant using the ELISA method, comparing the secretion levels in the engineered strain under dark conditions, continuous green light stimulation for 4 hours, and a “Switch” group with 2 hours of stimulation followed by 2 hours of light-off. (Data are presented as the mean ± SD. n=3. * 0.01<p<0.05, ** 0.001<p<0.01, *** p<0.001.)

Expressing sfGFP as reporter protein, the optogenetic function of the opto-effector EcN was rapidly activated within 2 hours, reaching 5.5 times of sfGFP expression higher than that before irradiation (**Fig 3B**). To shut off the optogenetic circuit, we found that after irradiating with green light for 1 hour and turning off the capsule LED followed by strain cultivation in a dark environment. The expression rate of sfGFP was also severely weakened within 1 hour (**Fig S5B**), demonstrating the time accuracy and light dose dependence of capsule-to-EcN optogenetic regulation. Considering the capsule was similar to a point light source with a limited radiation range, the increased distance of the light source from bacteria might reduce the activation of the cellular optogenetic function. When using capsules to give light signals for 2 hours to the opto-effector EcN in eight micro wells arranged side by side, the sfGFP expression intensities gradually decreased along with the increased distance and obstruction of light between each well (**Fig 3C and Fig S5C**). This result demonstrated the spatial accuracy of the capsule-to-EcN signal transmission function

The CcaS-CcaR-based opto-effector EcN showed high expression intensity after induction, stronger than commonly used strong expression systems in EcN (**Fig S6**). The CcaS-CcaR-based opto-effector EcN was also proved to be highly sensitive to green light. After gavage of the opto-effector EcN using LuxA as reporter (referred as opto-Lux EcN strain) to mice, the light emitted by the LED array only needed to be irradiated above the mouse cage (**Fig 3D and E**) to activate the engineered bacteria in the intestine within 2 hours (**Fig 3F**). Consistent with previous data (**Fig S5B**), after 2 hours of green light exposure to mice followed by removing of the light for 2 hours, a drastic reduction in the activated phenotype of opto-Lux EcN was observed similar the sfGFP reporter (**Fig S7**).

In order to employing the optogenetic controlled EcN for potential biomedical application, a scenario was devised to regulate the adhesion capacity of EcN via capsule-emitting light signals, wherein an opto-adhesion EcN strain carrying a genetic circuit for optogenetically controlled adhesive curli matrix secretion was constructed (**Fig 3G**). By editing the EcN genome to enhance the secretion of CsgA, and simultaneously regulating the secretion of CsgA-mfp3s fused with mussel foot mucin^23^ through optogenetics, bacterial adhesion to various substrates was enhanced. Crystal violet staining experiments showed that the strain’s adhesion to plastic microporous substrates was enhanced under capsule-to-EcN light control (**Fig 3H and I**).

Another scenario that often emphasized in biomedical applications we have implemented was to regulate protein secretion via capsule-to-EcN signal transmission. We introduced the hemolysin I secretion system gene HlyBD from uropathogenic *Escherichia coli* (UPEC) and used the CcaS-CcaR optogenetic tool to control the secretion of the protein fused with the secretion signaling peptide, constructing the protein-secreting opto-effector EcN strain. The concept of using EcN-to-capsule signaling system to transmit green light to control the ability of the engineered EcN to secrete functional proteins was tested and validated in following experiments (**Fig 3J**). The nanobody against human TNF-α, which was functionally similar to adalimumab or infliximab used in inflammatory bowel disease (IBD) treatment^24^, was selected as the protein of interest ready for optogenetic controlled secretion (referred to as Nb-TNF later). When Nb-TNF-secreting opto-effector EcN was stimulated with capsule-to-EcN light for 1-2 hours, the desired protein band was detected in the supernatant (**Fig 3K**). ELISA quantitative data indicated that continuous green light irradiation led to a sustained increase in Nb-TNF in the supernatant. The light exposure was terminated for 2 hours, leading to a slowdown in the subsequent increasing trend of the Nb-TNF concentration (**Fig 3L**). Consistent with previous data (**Fig S5B**), the expression and secretion of the nanobody showed time accuracy and green light dose dependence, enabling the therapeutic dose control by adjusting the duration of green light irradiation.

### Validation of EcN-to-capsule signal transmission *in vivo*

When applying the engineered EcN and the capsule in 50-90 kg pigs, the capability of EcN-to-capsule bioluminescent signal transmission was tested for monitoring the response of EcN status *in vivo*, especially the response of EcN to nitrate or arabinose molecules (**Fig 4A**). The capsule was placed into the intestine non-invasively by following approach. We used a moderately soft pipeline to assist placing the capsule inside the intestine; After turning on the voltage 1 and sensor button, the bioluminescence of the engineered EcN could be monitored. After the capsule was placed into the rectum for no less than 40 cm, it could still be connected via smartphone Bluetooth in a relatively broad space (radius of > 30 cm) near the pig’s posterior abdomen (**Fig 4B**). Due to the better light avoidance effect in the body, the dark photocurrent value measured by the capsule placed in the intestine was obviously lower than the corresponding value measured under the *in vitro* dark condition. The average blank value detected by the capsule in the body is only about 86.6 μA, which was subtracted as the base value in all *in vivo* experiments (**Fig S8**).

**Figure 4.**
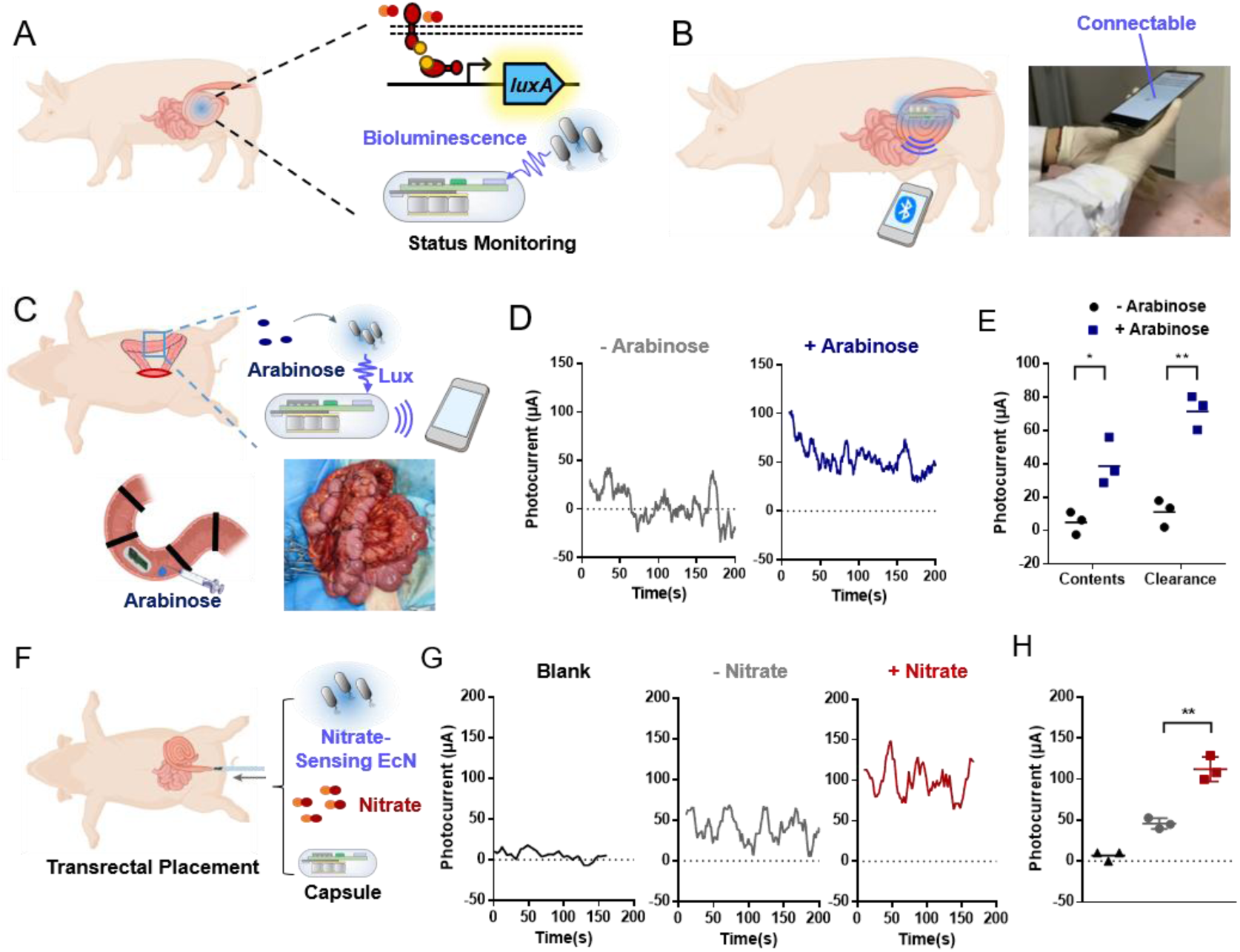
EcN-to-Capsule signal transmission *in vivo*. (A) The overview schematic of the *in vivo* EcN-to-capsule signal transmission for *in-situ* bacterial bioluminescence monitoring. **(B)** The image of the non-invasive placement of capsule through the anus in anesthetized pigs (left) and APP-controlled remotely received bioluminescent signals by capsule *in vivo* (right). **(C)** Schematic diagram of surgical placement of the capsule for bioluminescence monitoring (upper). Schematic diagram (bottom left) and photograph (bottom right) of performing multiple condition tests through tying the colon into several segments. **(D)** Typical bioluminescence signals of the ara-sensing EcN without (left) and with (right) arabinose addition in the colon. **(E)** Bioluminescence value of the ara-sensing EcN in response to arabinose in the environments of cleaned or un-cleaned intestinal contents. **(F)** Schematic diagram of the capsule monitoring the response of nitrate-sensing EcN to nitrate in the intestine. **(G)** Typical bioluminescence signals of different conditions: the blank background signals in the intestine without ara-sensing EcN existing (left), leakage bioluminescent signals of nitrate-sensing EcN without nitrate (middle), and the bioluminescent signals of nitrate-sensing EcN after adding exogenous nitrate (right). **(H)** Statistics of the bioluminescent signals in the blank intestine, intestinal nitrate-sensing EcN without nitrate addition, and intestinal nitrate-sensing EcN with exogenous nitrate addition. (Data are presented as the mean ± SD. n=3. * 0.01<p<0.05, ** 0.001<p<0.01, *** p<0.001.)

At first, to realistically simulate the process of monitoring the bioluminescent signals of the engineered EcN by using the capsule in the colon *in vivo*, the colon of anesthetized pigs was exposed through surgery for simulation experiments. The colon was tied into several segments by wrapping it with sutures to provide different operations, such as ara-sensing EcN injection, cleaning the intestinal contents or not, and arabinose doses. Capsule was surgically placed to monitoring the statuses in different segments (**Fig 4C**). Remotely monitoring the ara-sensing EcN by the capsule, the bioluminescence responded to arabinose could be significantly detected by the capsule (**Fig 4D**). Regardless of whether intestinal contents were cleared or not, the capsule could monitor the response of ara-sensing EcN to the inducer arabinose in the form of increased photocurrent, suggesting the effective *in vivo* application potential (**Fig 4E**). After cleaning the intestinal contents, the absolute intensity of bioluminescence signals detected in the form of photocurrent was even higher.

Furthermore, we investigated the ability of nitrate-sensing EcN to respond to nitrate in the intestine, and the EcN-to-capsule signal transmission functionality *in vivo* (**Fig 4F**). The nitrate-sensing EcN were infused into the pig intestine extensively and uniformly, and incubated for no less than 2 hours after adding nitrate solution. The capsule was placed in the colon by the non-invasive method to monitor the bioluminescence signal of the engineered EcN in response to nitrate. Although nitrate-sensing EcN exhibited a leakage expression signal compared to the blank signal, the bioluminescent signals in response to exogenous addition of nitrate visibly increased after 10 mM of nitrate addition (**Fig 4G**), and generated a significantly doubled signal enhancement (**Fig 4H**).

The above results indicate that the bioluminescent signals could effectively transmitted from engineered bacteria to electronic capsules in large animals, enabling the monitoring of the biomarker-responsive status of engineered bacteria in pigs. The above exploration provided a methodological basis for monitoring the state of engineered bacteria in large animals and more comprehensively evaluating the *in vivo* processes of engineered microbial therapeutics.

### Validation of capsule-to-EcN signal transmission *in vivo*

Next, the capability of capsule-to-EcN optogenetic signal transmission for regulating the function of engineered EcN within the intestine *in vivo* was evaluated. Opto-effector EcN and the electronic capsule were applied in live pigs weighting 50-90 kg. Transmission of signals and regulation of optogenetic function of opto-effector EcN was validated in the form of measurement of gene activated expression and secretion of the expressed proteins (**Fig 5A**). Following the placement of the electronic capsule into the rectum, it could be connected to and controlled *via* a smartphone to manage the LED on and off functions within the intestines (**Fig 5B and supplementary video 1**).

**Figure 5.**
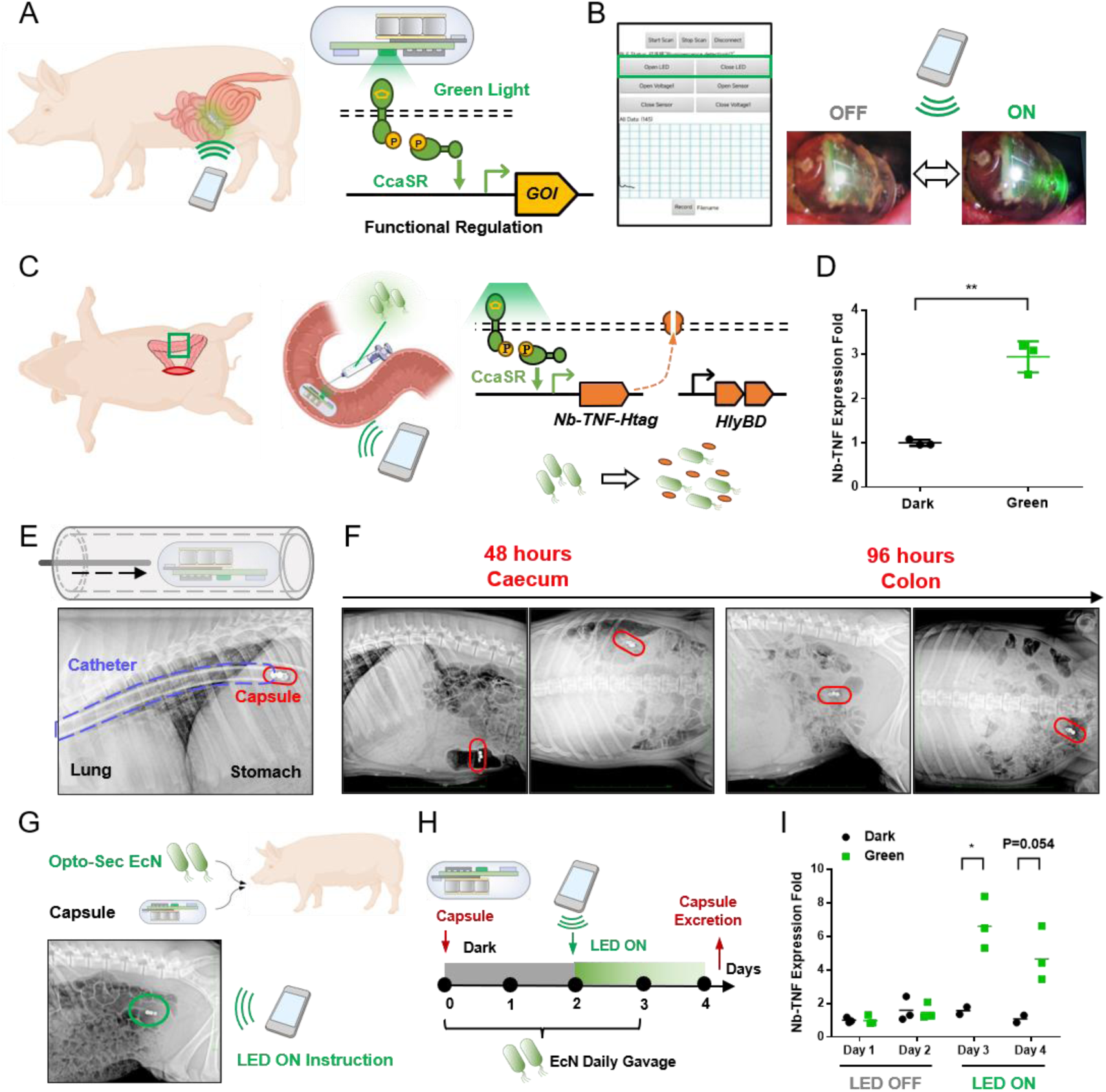
Capsule-to-EcN signal transmission *in vivo*. (A) The overview schematic of the *in vivo* capsule-to-EcN signal transmission for bacterial optogenetic functional regulation. **(B)** The remote control of capsule LED *in vivo* by APP (upper) and the LED switch images under colonoscopy (bottom). **(C)** Schematic diagram of the surgical placement of opto-effector EcN and capsule for optogenetic regulation of EcN. **(D)** The expression of Nb-TNF under optogenetic control of green light of capsule LED for 2 hours (group “Green”) compared with capsule LED off for 2 hours (group “Dark”). **(E)** The process of placing the electronic capsule into the esophagus via a gastric catheter (X-ray image). **(F)** The process of electronic capsule passing through the whole digestive tract (X-ray images in every two days). **(G)** Schematic diagram of the experiment of orally administrating opto-effector EcN and capsule for optogenetic regulation of EcN. **(H)** Timeline diagram of EcN and capsule administration and capsule LED open instruction. **(I)** The expression of Nb-TNF under optogenetic control of green light of capsule LED, during which the “Green” group was received capsule LED open instruction after Day 2, while the “Dark” group was kept LED off in the whole process. (Data are presented as the mean ± SD. n=3. * 0.01<p<0.05, ** 0.001<p<0.01, *** p<0.001.)

At first, to realistically simulate the process of the capsule regulating opto-effector EcN in the colon *in vivo* while facilitating the sampling analysis for the secretion of the target protein, the capsule was surgically placed into the colon of anesthetized pigs. At the same time, opto-effector EcN with a CFU of 10^9^ was injected near the capsule (**Fig 5C**). The colon was then placed back to the pig body. Under remotely control, the capsule regulated the expression of Nb-TNF by the engineered bacteria at the localized intestinal segment. Sampling of the intestinal contents to detect the expression of Nb-TNF mRNA showed that after 2 hours of light stimulation by the electronic capsule, the opto-effector EcN effectively expressed more Nb-TNF proteins (**Fig 5D**).

Next, the opto-effector EcN and capsule were orally administrated to evaluate their whole passage in the digestive tract. The capsule was able to maintain the photosensor at “ON” state for about 10 minutes, or maintain the LED green light at “ON” state for more than 36 hours (**Fig S7**), or keep the Bluetooth connection for more than two weeks in a standby state. Therefore, for the *in vivo* application of optogenetic regulation, the capsule’s endurance time was sufficient. As for the oral administration of electronic capsule in pig, for avoiding the damage on capsule by mouth chewing force, the capsule was carried through a gastric catheter and inserted into the pig’s esophagus, and then assisted by a rod to be push deeply into the pig’s esophagus (**Fig 5E and S9A**). It took approximately 5 days for the capsule to pass through the digestive tract in 50-90 kg pigs (**Fig 5F**), maintaining connectable throughout the days (**Fig S9B and supplementary video 2**). The Bluetooth signal intensity for the capsule *in vivo* fluctuated with its location due to tissue depth, yet remained above the connectivity threshold (>100 dB) even at its deepest within the body (**Fig S9C and D**).

By remotely controlling capsule LED through a smartphone, the expression of opto-effector EcN was activated (**Fig 5G**). The opto-effector EcN were daily orally administrated; After 48 hours of orally administrated of capsule, EcN were activated by continuous LED light for the rest time (**Fig 5H**). Daily fecal bacterial mRNA analysis showed that Nb-TNF gene expression specifically increased in the Day 3 and 4 when LED light was provided for more than 24 hours (**Fig 5I**).

The above results indicated that the LED light from the electronic capsule effectively activated the predesigned functionality of the engineered bacteria in pigs through regulation of the cellular optogenetic circuits. The above exploration provided a methodological basis for future studies on more complex optogenetic function regulation using large animal models, and established a convenient method to remotely controlling the engineered microbial therapeutics.

### The monitoring-regulating cycle based on bidirectional bio-electronic optical communication

On the basis of the above result, it had been possible to establish a monitoring-regulating cycle with the engineered EcN *in vitro* and *in vivo* via bidirectional optical language between the capsule and engineered EcN community. In this monitoring-regulating cycle, a first sensor EcN strain transmitted bioluminescent signals to the capsule to detect environmental molecules of the bacterial community, and a second effector EcN strain received optogenetic signals from the capsule to regulate the function of the bacterial community (**Fig 6A**). Therefore, it was possible to monitor the response to the environment *in situ* and regulate bacterial function accordingly.

**Figure 6.**
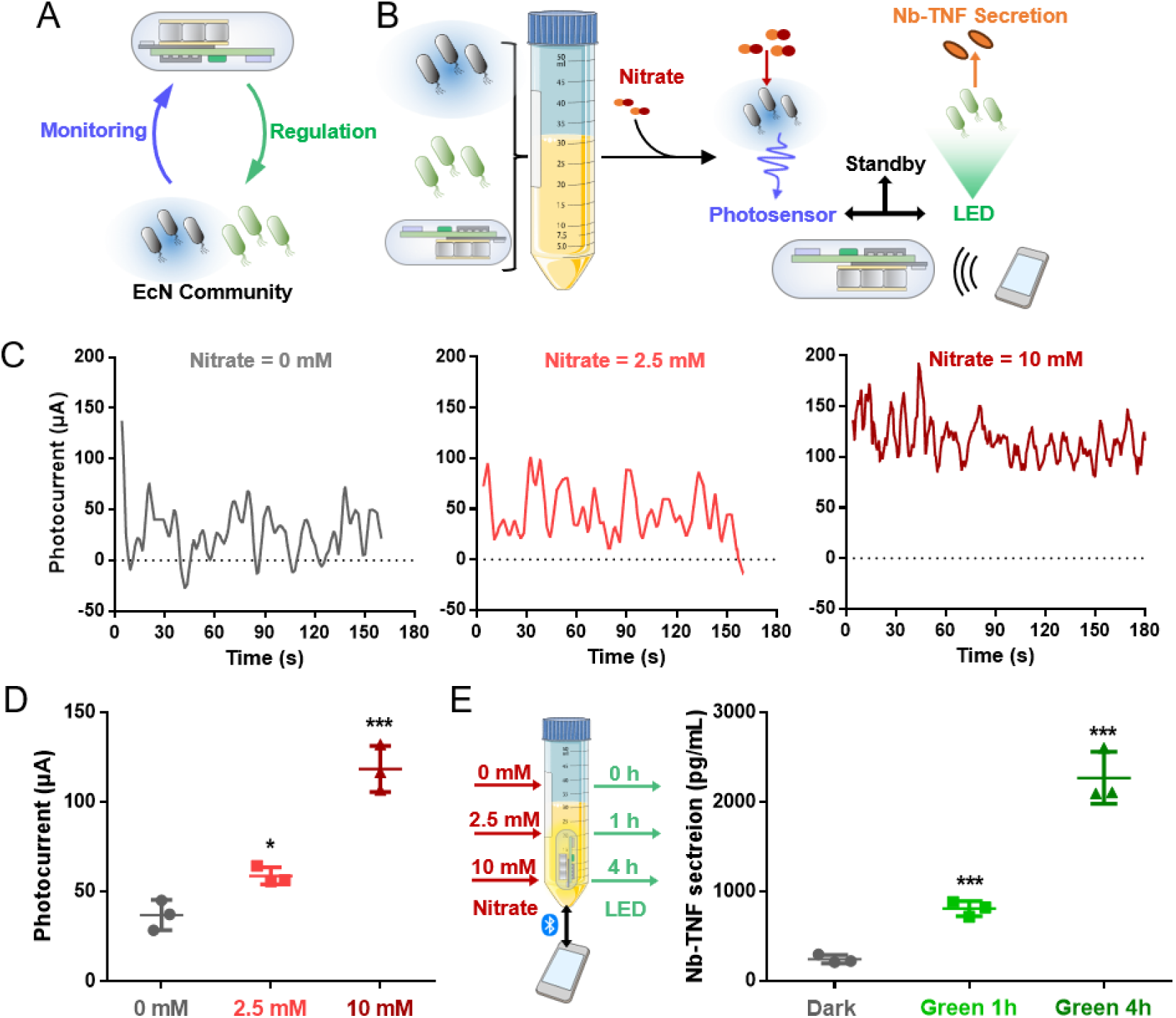
Establishing monitoring-regulating cycle of engineered EcN community. (A) The overview schematic of the monitoring-regulating cycle based on bidirectional bio-electronic optical communication. **(B)** The schematic diagram of the trail of monitoring and regulating engineered EcN community by controlling capsule functions. **(C)** Typical bioluminescence signals of engineered EcN community exposed in different nitrate conditions (0, 2.5, 10 mM). **(D)** Statistics of the bioluminescent signals of engineered EcN community exposed in different nitrate conditions. **(E)** Schematic diagram of the trail (left) and concentration of Nb-TNF secretion (right) of optogenetic functional control of the engineered EcN community based on pre-monitored bioluminescence.

A simulative trail was conducted *in vitro*, in which nitrate-sensing EcN strain and Nb-TNF-secreting EcN strain were co-cultured with liquid culture medium in a 50 ml centrifuge tube (**Fig 6B**). The tube was kept on standby mode until in demand. After nitrate was added in different concentration for 2 hours, the photosensor of the capsule was switched on to monitor the EcN response. When the monitoring result was positive, accordingly, the LED of the capsule was switched on for a different time period to control the secretion of Nb-TNF. The experiment result showed that the EcN-to-capsule bioluminescent signal monitoring function achieved significant difference of photocurrent in response to the various nitrate conditions (**Fig 6C and D**). According to the bioluminescent signals of 0, 2.5, and 10 mM of nitrate, we chose 0, 1, and 4 hours of optogenetic signals to stimulate Nb-TNF secretion. As a result, the Nb-TNF secreted to the supernatant was dose-relied according to the concentration of nitrate (**Fig 6E**).

The monitoring-regulating cycle based on bidirectional bio-electronic optical communication provided a potential strategy for the integrated diagnose and treatment of intestinal diseases. It enabled the detection of the biomarkers in the host’s intestine, and regulation of the therapeutic dose of engineered bacterial function remotely in real time. There have reported several synthetic biological diagnostic and therapeutic strategies directly implemented through genetic circuits in the engineered strain, for example, to express genes playing memory, reporter, and therapeutic functions under the control of IBD biomarker thiosulfate^4^, and to express of *Pseudomonas aeruginosa* killing genes under the control of their quorum sensing signals^25^. Different from the above strategies of autonomous engineered bacteria, our strategy depended on the motivated involvement of human operator and the decision made by human operator to sophisticatedly regulate therapeutic doses, which might be irreplaceable in clinical practice.

## Discussion

Synthetic biology technologies enable engineered microorganisms becoming a promising new type of live biotherapeutic products. However, the lack of methodology for *in situ* monitoring of the status of engineered microbes and real-time regulation of their functions results in insufficient understanding and control of their *in vivo* functions.

In this work, through collaborative design between the engineered bacteria EcN and electronic capsules, we established the bi-directional communication between human operator and intestinal microbes via optical language. The bioluminescent module was composed of a genomic integrated *lux-BCDE* gene cluster and a separated *LuxA* participating in functional update in the genetic circuits, making it flexible to be a reporter gene. Moreover, its luminous intensity was high, allowing easy penetration through absorbent media such as chyme and feces in the intestine. The optogenetic module based on CcaS-CcaR was one of the highest inducible two-component optogenetic tools reported so far^21^, applied in the strains constructed in this work, it demonstrated sensitive light regulation ability and extremely strong functional gene expression.

In both *in vitro* and *in vivo* trails, the electronic capsules could receive bioluminescence signals from the engineered EcN, and then translate it into a language of wireless signal and transmitted it in a real-time form to the human operator’s smartphone, which was the function of microbial monitoring. Then, under the operator’s control, a new wireless signal would turn on the LED in capsule and emit light to regulate the engineered strains, which was the function of microbial stimulating. This process is a microcosm of the entire process of medical personnel diagnosing intestinal health, making decisions, and intervening diseases. By contrast, there are already various diagnostic and therapeutic strategies for intestinal diseases reported, such as the engineered bacteria that directly activate therapeutic genes in response to biomarkers^4,25,26^, implantation systems for responsive drug release^27–29^, etc., they behave in a closed-loop manner autonomously *in vivo* without solving the problem of in situ diagnosis cannot be real-time monitored and uncontrollable treatment.

Our work provides a methodological interface for the study of intestinal engineered bacteria at the animal level. This article aims to demonstrate the feasibility of using light signals as a medium for communication between engineering bacteria and the outside world. The bioluminescence and optogenetics modules constructed in this article can also be extended to more functions, and are compatible with existing engineered bacteria designs that function in a closed-loop manner autonomously. By introducing ingestible microelectronic devices, our strategy exposes the diagnosis and treatment process *in vivo* to the supervision and control of users, which has irreplaceable clinical significance.

The outstanding works of Timothy K. Lu’s team in developing ingestible capsules that encapsulating engineered bacteria as sensors^15,16^ has given us confidence in our strategy. Unlike the fundamental purpose of their works, we regard engineered bacteria as the core function of intestinal diagnosis and treatment, and the electronic capsules are introduced only as crossroads of information. It enables more flexible application of massive engineered bacteria, making it possible to use engineered bacteria as therapeutic agents. At the same time, it allows for the inclusion of a wider range of monitoring purposes, such as monitoring the quantity distribution and functional activity of therapeutic engineered bacteria.

As a methodological study on the application of engineered live biotherapeutic products, there are still many aspects worth optimizing and improving in our strategy. Firstly, the engineered bacteria designed in this work did not ultimately meet the clinical translational needs of disease diagnosis and treatment. Before conducting researches for clinical purposes, the design of engineered bacteria should be plasmidless and have no antibiotic resistance genes. Similarly, there is space for optimization and compression of the size and energy consumption of electronic capsules. Finally, there was only one signal dimension that could be monitored in this research, which limited the potential of extending it to simultaneous detection of more indicators.

In future research, there are several strategies for collaborative design of engineered bacteria and capsules that can expand the dimension of bacteria-to-capsule signals. For example, it is envisioned that by introducing an orthogonal luciferase-luciferin tools^30^, capsules can release luciferin on demand and sequentially detect multiple signals from bacteria. Alternatively, orthogonal quorum sensing genes^31^ can be designed for engineered bacteria as output signals, applying bacteria-encapsulating capsules design multiple engineered bacterial sensor channels that respond to each quorum sensing molecules. In summary, our strategy can be compatible with increasingly complex and robust synthetic biological designs and microelectronic designs, thus having increasingly strong practical potential.

## Methods

### Strains and culture conditions

All plasmids and bacterial strains used in this study were listed in Supplementary Table 1 and Table 2. The strain *Escherichia coli* DH5α were used for plasmid construction and cloning. The strain *E. coli* Nissle 1917 were used as the chassis for genomic edition and transformation the plasmids for the fully functional genetic circuits.

Luria-Bertani (LB) broth and plates were used for strain selection and propagation. Kanamycin (50 mg L^-1^), ampicillin (100 mg L^-1^), and / or chloramphenicol (50 mg L^-1^) was added appropriately according to different antibiotic resistance conditions.

For the experiments of activating protein secretion by optogenetic control, M9 medium was used, with a final concentration of 2 mM MgSO_4_, 0.1 mM CaCl_2_, and 0.4% glucose supplemented. Kanamycin (50 mg L^-1^), ampicillin (100 mg L^-1^), and / or chloramphenicol (50 mg L^-1^) was also added according to conditions.

### Plasmids and strains construction

All plasmids were constructed by seamless clone using 2X MultiF Seamless Assembly Mix (RK21020, ABclonal Technology Co.,Ltd., China). DNA fragments were amplified using PCR using PrimeSTAR® Max DNA Polymerase (Takara Biomedical Technology (Beijing) Co., Ltd., China). Genes such as *ccaS*, *ccaR*,^21^ *Nb-TNF*^32^ were synthesized in Azenta Co. Ltd., China, based on reported sequences. HlyB and HlyD were amplified from uropathogenic *Escherichia coli*. Genomic modified strains EcN-CSG and EcN-Lux were constructed based on the optimized CRISPR/Cas9-assisted recombineering method^33^.

### Preparation and application of electronic capsules

Before using the electronic capsule, a group of new button cells was welded to the customized printed circuit board and the shell was molded. The Bluetooth connection with the capsule and the control of the LED or photoelectric sensor are completed through a self-designed app. When testing the functional regulation and status monitoring of the engineered bacterial strain based on the electronic capsule, the strain is cultured and the electronic capsule is placed inside it, and its LED is controlled wirelessly to regulate its photogenetic function, or its photoelectric sensor is controlled to monitor the bioluminescence signal from the engineered bacteria. Since the photoelectric sensor of the electronic capsule is suitable for monitoring very weak bioluminescence, when using this function in vitro, the capsule should be placed in a strict light-avoiding environment.

### Characterization of genetic circuits

For bioluminescent strains, 1% of the bacteria cultured overnight under uninduced conditions were re-cultured for the preparation of tests. In order to use constitutive bioluminescent bacteria for the basic quantification, the gradient dilution of the constitutive expression bacterial solution with OD 600 being approximately 0.5 was used for microplate reader or capsule detection. In order to testify the function of further genetic circuits using bioluminescent reporter, induce the bacterial solution in the result with different concentration and time according to the experimental setting.

For optogenetic strains, transformation and cultivation should be carried out in a dark environment that avoids any direct light exposure. 1% of the bacteria cultured overnight under dark conditions were re-cultured for the preparation of light stimulation. When the OD 600 was 0.3-0.5, the bacteria were transferred to the corresponding light induced environment to testify gene expression.

In order to accurately detect the secretion of the target product of bacteria, the bacteria used in such experiments were cultured in M9 medium. The bacterial solution was collected of the light induced group, dark control group, or constitutive starter strains cultured for a certain period of time after inoculation. Centrifuge at 10000 rpm and filter through a 0.22 μm pore size filter sequentially. Then the fluorescence values were measured by microplate reader. Western Blot and Enzyme-Linked ImmunoSorbent Assay (ELISA) based on HA-tag antibody were carried out to validate protein secretion in the bacterial culture supernatant.

### *In vitro* simulation trails using intestinal tissues

The intestinal tissue of pigs was used for in vitro simulation experiments. During the experiment, a corresponding strain of approximately 1mL was uniformly adhered to the inner wall of approximately 20 cm long intestinal tissue after cleaning. Afterwards, use a syringe to locally inject 100uL of arabinose or potassium nitrate solution into the center of the intestinal tissue. After incubating at 37 ° C for 2 hours, manually insert the capsule from one end of the intestinal tissue in a dark environment and push it at a constant speed. During this period, control the electronic capsule to maintain reading and recording of photoelectric signals. Finally, in order to visually demonstrate the activation effect of engineering bacteria after local injection of arabinose or potassium nitrate, the intestinal tissue was cut along the busbar and its inner wall was exposed, and its bioluminescence was imaged using a bioluminescence imaging system.

### Mouse experiments

Male C57BL/6 mice (7 weeks) were purchased from SPF (Beijing) Biotechnology Co., Ltd. Animal experiments were carried out after the mice were reared adaptively for one week. Animal experiments followed the requirements of the People’s Republic of China (GB14925–2010).

5*10^8^ of bacterial count of engineered EcN were collected and dispersed into 200 μL of saline for mice gavage. After the engineering bacteria were gavaged and stabilized for 2 hours, 100 uL of arabinose, potassium nitrate solution, or physiological saline as a control were gavaged. Within 2-4 hours after that, the distribution of bioluminescence in the abdomen was periodically observed using a bioluminescence imaging system. When inducing optogenetic bacteria in the intestinal of mice, hair removal was performed on the abdomen of the mice using hair removal cream in advance. The mice were anesthetized with isoflurane and then fixed on their backs. Within 2-4 hours after that, the distribution of bioluminescence in the abdomen was periodically observed using a bioluminescence imaging system.

### Pig experiments

Quarantined healthy ordinary domestic pigs (castrated males) weighting 50-90kg were used for live pig experiments. Animal experiments were carried out after the pigs were reared adaptively for one week. Animal experiments followed the requirements of the People’s Republic of China (GB14925–2010).

In order to non-invasively place optical electronic capsule devices into the body to study their functions, two methods are used in this chapter: oral delivery of capsules and transanal placement of capsules. Mix a small amount of feed (not exceeding 200g) with a dosage of CFU=10^11^ of engineering bacteria to prompt the pigs to quickly ingest the engineered bacteria. Feed the pigs with engineered bacteria daily.

For the former, after injecting the pig with the muscle, the electronic capsule was pushed into the esophagus with the help of a gastric tube, and the capsule was connected to the smartphone every day for the next 4 days to complete the subsequent experimental operation. In order to monitor the process of the capsule being transported in the gastrointestinal tract, after delivering the capsule into the esophagus orally, at specific time intervals, it was anesthetized with Zoletil (5mg/kg) and xylazine hydrochloride (2mg/kg), and its location was photographed using an animal X-ray imaging system.

For the latter, the pigs were fasted for 24 hours and deprived of water for 4 hours, and underwent lactulose enema for two hours before surgery. After emptying the intestines, the pigs were anesthetized with intramuscular injection of Zoletil (5mg/kg) + xylazine hydrochloride (2mg/kg), and the electronic capsule was temporarily placed in the colorectal tract using an endoscope to complete subsequent experimental procedures. After placing the capsule in the intestine by any means, the capsule signals were remotely searched through the APP, and the location near the capsule is found to complete the Bluetooth connection with the capsule. The function of the LED or photoelectric sensor can be remotely controlled.

After the experiment, 10% KCl solution (0.5 mL/kg) was rapidly injected intravenously into the pig under anesthesia and euthanized.

## Acknowledgments

We thank the Laboratory Center of the School of Life Sciences (Tianjin University) and experimental animal center of Northwest Agriculture and Forestry University for facility support. This work was supported by the National Key Research and Development Program of China (2019YFA0906500), the National Natural Science Foundation of China for Excellent Young Scholars (32122047), Tianjin Natural Science Foundation for Distinguished Young Scholars (23JCJQJC00210), Beijing-Tianjin-Hebei Basic Research Cooperation Project of Beijing Natural Science Foundation (23JCZXJC00370), and the Key Program of Tianjin Natural Science Foundation (22JCZDJC00230).

## Notes

### Competing Interest Statement

The authors have declared no competing interest.

## References

1 Riglar, D. T. & Silver, P. A. Engineering bacteria for diagnostic and therapeutic applications. Nat Rev Microbiol 16, 214–225, doi:10.1038/nrmicro.2017.172 (2018).

2 Cubillos-Ruiz, A. et al. Engineering living therapeutics with synthetic biology. Nature Reviews Drug Discovery 20, 941–960, doi:10.1038/s41573-021-00285-3 (2021).

3 Scott, B. M. et al. Self-tunable engineered yeast probiotics for the treatment of inflammatory bowel disease. Nature medicine 27, 1212–1222, doi:10.1038/s41591-021-01390-x (2021).

4 Zou, Z. P., Du, Y., Fang, T. T., Zhou, Y. & Ye, B. C. Biomarker-responsive engineered probiotic diagnoses, records, and ameliorates inflammatory bowel disease in mice. Cell Host Microbe 31, 199–212 e195, doi:10.1016/j.chom.2022.12.004 (2023).

5 Luke, J. J. et al. Phase I Study of SYNB1891, an Engineered E. coli Nissle Strain Expressing STING Agonist, with and without Atezolizumab in Advanced Malignancies. Clinical cancer research : an official journal of the American Association for Cancer Research 29, 2435–2444, doi:10.1158/1078-0432.Ccr-23-0118 (2023).

6 Vockley, J. et al. Efficacy and safety of a synthetic biotic for treatment of phenylketonuria: a phase 2 clinical trial. Nature metabolism 5, 1685–1690, doi:10.1038/s42255-023-00897-6 (2023).

7 Puurunen, M. K. et al. Safety and pharmacodynamics of an engineered E. coli Nissle for the treatment of phenylketonuria: a first-in-human phase 1/2a study. Nature metabolism 3, 1125–1132, doi:10.1038/s42255-021-00430-7 (2021).

8 Kurtz, C. B. et al. An engineered E. coli Nissle improves hyperammonemia and survival in mice and shows dose-dependent exposure in healthy humans. Science translational medicine 11, doi:10.1126/scitranslmed.aau7975 (2019).

9 Martínez, E. P. & Robles, E. P. Capsule endoscopy and deep enteroscopy. Endoscopy 46, 787–790, doi:10.1055/s-0034-1377448 (2014).

10 Ding, Z. et al. Novel scheme for non-invasive gut bioinformation acquisition with a magnetically controlled sampling capsule endoscope. Gut 70, 2297–2306, doi:10.1136/gutjnl-2020-322465 (2021).

11 Hou, B. et al. A swallowable X-ray dosimeter for the real-time monitoring of radiotherapy. Nature Biomedical Engineering 7, 1242–1251, doi:10.1038/s41551-023-01024-2 (2023).

12 Kalantar-Zadeh, K. et al. A human pilot trial of ingestible electronic capsules capable of sensing different gases in the gut. Nature Electronics 1, 79–87, doi:10.1038/s41928-017-0004-x (2018).

13 Ramadi, K. B. et al. Bioinspired, ingestible electroceutical capsules for hunger-regulating hormone modulation. Science robotics 8, eade9676, doi:10.1126/scirobotics.ade9676 (2023).

14 Srinivasan, S. S. et al. RoboCap: Robotic mucus-clearing capsule for enhanced drug delivery in the gastrointestinal tract. Science robotics 7, eabp9066, doi:10.1126/scirobotics.abp9066 (2022).

15 Inda-Webb, M. E. et al. Sub-1.4 cm(3) capsule for detecting labile inflammatory biomarkers in situ. Nature 620, 386–392, doi:10.1038/s41586-023-06369-x (2023).

16 Mimee, M. et al. An ingestible bacterial-electronic system to monitor gastrointestinal health. *Science (New York*, N.Y.) 360, 915–918, doi:10.1126/science.aas9315 (2018).

17 Kim, C. K., Adhikari, A. & Deisseroth, K. Integration of optogenetics with complementary methodologies in systems neuroscience. Nature reviews. Neuroscience 18, 222–235, doi:10.1038/nrn.2017.15 (2017).

18 Nihongaki, Y., Kawano, F., Nakajima, T. & Sato, M. Photoactivatable CRISPR-Cas9 for optogenetic genome editing. Nature biotechnology 33, 755–760, doi:10.1038/nbt.3245 (2015).

19 Azad, T. et al. Luciferase-Based Biosensors in the Era of the COVID-19 Pandemic. ACS nanoscience Au 1, 15–37, doi:10.1021/acsnanoscienceau.1c00009 (2021).

20 Hoffman, S. M., Tang, A. Y. & Avalos, J. L. Optogenetics Illuminates Applications in Microbial Engineering. Annual review of chemical and biomolecular engineering 13, 373–403, doi:10.1146/annurev-chembioeng-092120-092340 (2022).

21 Ong, N. T. & Tabor, J. J. A Miniaturized Escherichia coli Green Light Sensor with High Dynamic Range. Chembiochem 19, 1255–1258, doi:10.1002/cbic.201800007 (2018).

22 Hartsough, L. A. et al. Optogenetic control of gut bacterial metabolism to promote longevity. eLife 9, doi:10.7554/eLife.56849 (2020).

23 An, B. et al. Programming Living Glue Systems to Perform Autonomous Mechanical Repairs. Matter 3, 2080–2092, doi:10.1016/j.matt.2020.09.006 (2020).

24 Cosnes, J. et al. Adalimumab or infliximab as monotherapy, or in combination with an immunomodulator, in the treatment of Crohn’s disease. Alimentary pharmacology & therapeutics 44, 1102–1113, doi:10.1111/apt.13808 (2016).

25 Hwang, I. Y. et al. Engineered probiotic Escherichia coli can eliminate and prevent Pseudomonas aeruginosa gut infection in animal models. Nat Commun 8, 15028, doi:10.1038/ncomms15028 (2017).

26 Mao, N., Cubillos-Ruiz, A., Cameron, D. E. & Collins, J. J. Probiotic strains detect and suppress cholera in mice. Science translational medicine 10, doi:10.1126/scitranslmed.aao2586 (2018).

27 Zhao, P. et al. Nanoparticle-assembled bioadhesive coacervate coating with prolonged gastrointestinal retention for inflammatory bowel disease therapy. Nat Commun 12, 7162, doi:10.1038/s41467-021-27463-6 (2021).

28 Ghosh, A. et al. Gastrointestinal-resident, shape-changing microdevices extend drug release in vivo. Science advances 6, doi:10.1126/sciadv.abb4133 (2020).

29 Liu, J. et al. Triggerable tough hydrogels for gastric resident dosage forms. Nature Communications 8, 124, doi:10.1038/s41467-017-00144-z (2017).

30 Williams, S. J. et al. Biochemical Analysis Leads to Improved Orthogonal Bioluminescent Tools. Chembiochem 24, e202200726, doi:10.1002/cbic.202200726 (2023).

31 Du, P. et al. De novo design of an intercellular signaling toolbox for multi-channel cell-cell communication and biological computation. Nat Commun 11, 4226, doi:10.1038/s41467-020-17993-w (2020).

32 Lynch, J. P. et al. Engineered Escherichia coli for the in situ secretion of therapeutic nanobodies in the gut. Cell Host & Microbe 31, 634–649.e638, doi:10.1016/j.chom.2023.03.007 (2023).

33 Huang, C., Guo, L., Wang, J., Wang, N. & Huo, Y. X. Efficient long fragment editing technique enables large-scale and scarless bacterial genome engineering. Appl Microbiol Biotechnol 104, 7943–7956, doi:10.1007/s00253-020-10819-1 (2020).

